# Sound-evoked facial motion in ferrets: evidence for species differences in sensorimotor coupling

**DOI:** 10.64898/2026.03.09.710529

**Authors:** Mathilde Martin, Yves Boubenec

## Abstract

Sensory stimuli elicit uninstructed movements in the mouse, and cortical activity covaries with these movements, raising the question of how much apparent sensory encoding reflects behaviorally driven signals. Still, whether sounds systematically elicit uninstructed movements in non-rodent species is an open question. Here we characterize sound-evoked facial motion and pupil dynamics in head-fixed ferrets, a carnivore species, passively exposed to broadband, natural, and synthetic auditory stimuli. Ferrets reliably produced facial responses to sounds, but unlike mice, these responses were dominated by a single onset-locked component scaling with sound loudness, tracked only temporal modulations below 2 Hz, and carried no information about sound identity or category. Paradoxically, synthetic sounds, which preserved spectrotemporal statistics but lacked higher-order natural structure, elicited stronger responses than their natural counterparts for several sound categories, suggesting that ferret behavioral responses reflect selective sensitivity to acoustic novelty rather than ecological salience. Pupil dynamics mirrored facial motion, consistent with a shared arousal mechanism. Together, these results show that sound-evoked facial motion in ferrets is less stimulus-specific than in mice, and suggest that the tight sensorimotor integration observed in rodents may reflect a species-specific organization rather than a general mammalian principle.

## Introduction

Neural recordings in auditory cortex are typically interpreted as reflecting sensory encoding. Yet a growing body of evidence from rodents has revealed that a substantial fraction of trial-to-trial variability in cortical responses is driven not by the stimulus, but by concurrent uninstructed movements (Musall et al., 2019; Stringer et al., 2019; Yin et al., 2025). In mice, even during passive listening under head fixation, sounds reliably elicit facial movements, such as whisker pad deflections, snout motion, and eye movements (Clayton et al., 2024; Meyer et al., 2018), that covary with arousal state (Niell & Stryker, 2010; Parker et al., 2020). These movement-evoked signals are not limited to startle responses: they can be stimulus-locked, sound-specific and sensitive to low sound levels (Clayton et al., 2024; Yeomans et al., 2002). Critically, such facial movements predict widespread cortical activity across sensory areas in mice (Bimbard et al., 2023; Musall et al., 2019). This raises a fundamental interpretational question: how much of what we as auditory neuroscientists characterize as sensory encoding reflects, in part, a behaviorally-driven component (Olsen & Hasenstaub, 2025)?

The answer may be strongly species-dependent. In primates, spontaneous movements contribute only weakly to evoked cortical signals (Talluri et al., 2023), suggesting that the tight coupling between sound, movement, and cortical activity observed in mice does not necessarily generalize across mammals. Understanding where ferrets fall along this axis matters considerably for auditory neuroscience, because ferrets are an important non-rodent model organism in the field. Their larger, gyrified cortex and well-characterized auditory hierarchy, spanning primary and non-primary fields with processing properties that are routinely compared to those of humans, have made them a key bridge species for understanding auditory cortical computation (Landemard et al., 2021, 2025; Sabat et al., 2025). If ferret auditory cortex is as susceptible to behavioral confounds as mouse cortex, interpreting sound-evoked cortical activity could require accounting for behavioral contributions. If it is not, this would suggest that the tight sound-movement coupling observed in mice may reflect something idiosyncratic about rodent sensorimotor organization rather than a general feature of mammalian auditory processing, and that ferrets may represent a more tractable model for isolating sensory from behavioral contributions to neural activity. Yet it is entirely unknown whether passive sounds systematically elicit uninstructed facial movements in ferrets, and whether such responses are modulated by stimulus properties.

To address this question, we provide the first systematic characterization of sound-evoked facial motion and pupil dynamics in head-fixed ferrets during passive listening. Using high-frame-rate video and facial motion energy analysis, we tested responses to broadband stimuli with controlled temporal structure, natural sounds spanning multiple ecological categories, and spectrotemporally matched synthetic counterparts that preserved low-level statistics while lacking higher-order natural structure. We found that ferrets exhibited robust, onset-dominated facial responses to broadband sounds, that response amplitude scaled with loudness but did not depend on sound category, and that facial motion was stronger for some synthetic sounds than for their natural counterparts despite matched spectrotemporal statistics. These results demonstrate that sound-evoked facial motion in ferrets is sensitive to global acoustic features (sound level and sound naturalness), but does not reflect fine sound structure beyond spectrotemporal statistics, contrary to what is observed in mice.

## Results

### Broadband sounds elicit robust facial motion responses in ferrets

Broadband sounds elicited robust, time-locked facial motion responses in ferrets. We quantified facial motion energy (FME) in seven head-fixed female ferrets during passive listening to two types of broadband stimuli: repeated descending frequency sweeps and silent gaps in continuous white noise (Fig. 1A). For repeated sweeps (6 sweeps per trial; 1-4 Hz), FME showed an evoked response at sound onset at all rates (Fig. 1B-C, sweep-evoked response: paired t-test: *p* = 0.02). However, synchronisation to individual sweeps was visible only at the lowest rate (Fig. 1B bottom). Consistently, we found that synchronisation power, which quantified entrainment of the FME to the stimulus, decreased monotonically with rate (Fig. 1D, effect of sweep rate on synchronisation: repeated measures ANOVA: p< 10^*−*2^). At 1 Hz, the response peaked approximately 300 ms after sweep onset (Fig. 1C). For silent gaps, FME increased at noise onset following long gaps (0.2 and 0.5 s in Fig. 1E), with response latencies that were consistent with the latency observed in the responses to individual sweeps (Fig. 1C). Across both paradigms, enhanced synchronization at lower sweep rates and larger responses for the longest gaps suggest that sound-evoked facial motion in ferrets primarily tracks low-frequency temporal features (Fig. 1B-E). This pattern was confirmed by analyzing the high-dimensional representation of facial signal through principal component analysis, in which the first principal component captured the low-frequency temporal features of the sound (SupFig. 1-2).

**Figure 1.**
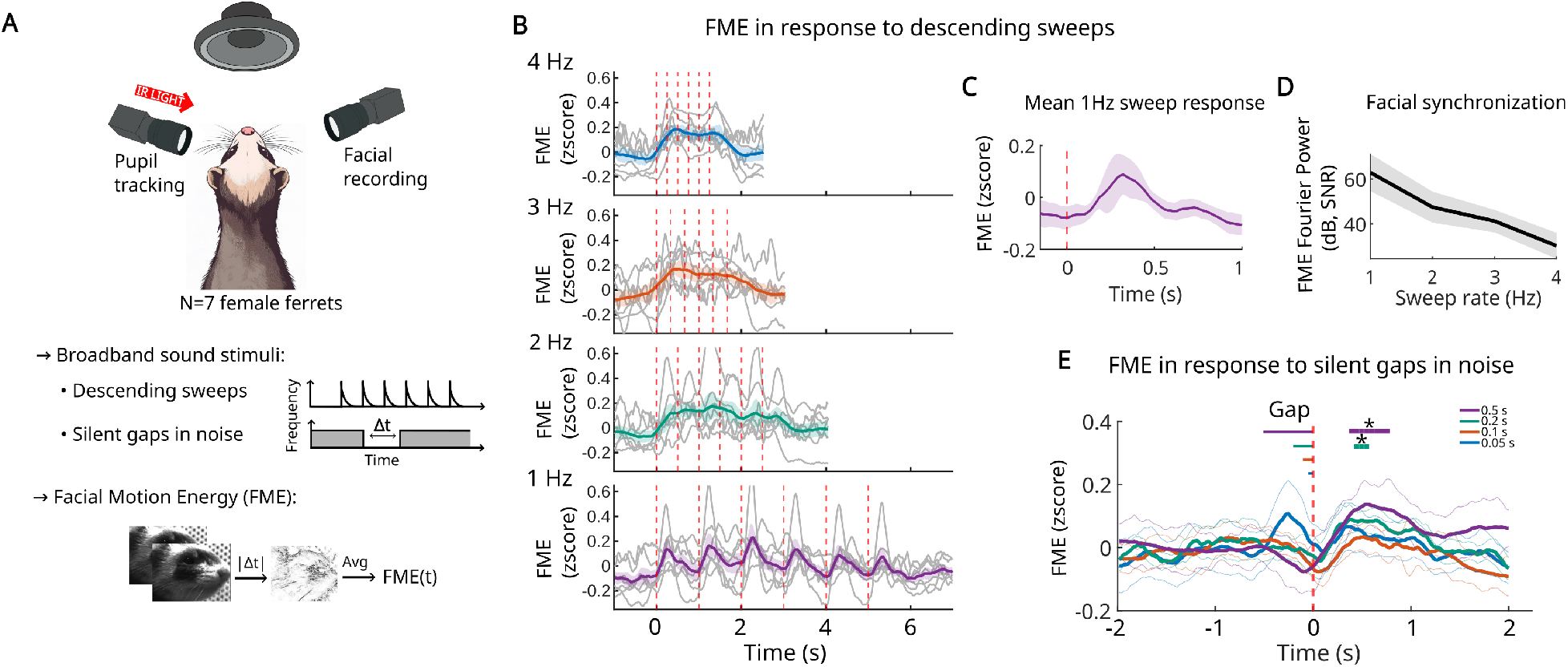
Broadband sounds elicit robust facial motion responses in ferrets. **A.** Experimental design and computation of Facial Motion Energy (FME). **B**. FME responses to descending sweeps at four presentation rates (1-4 Hz) (colored: across-subject mean; gray: individual subjects). Vertical dotted lines indicate sweep onsets. Sweep effect tested with a t-test on the mean response to the first two sweeps (*p* = 0.02 for each presentation rate, Benjamini-Hochberg corrected). **C**. FME response to descending sweeps presented at 1 Hz, averaged across subjects and sweeps. **D**. Facial synchronization as a function of sweep rate, quantified as Fourier power at each sweep frequency (dB relative to the high-frequency noise floor). Repeated measures ANOVA: *p* < 10^*−*2^. **E**. FME responses for different silent-gap durations, time-locked to gap offset (colored: across-subject mean; gray: individual subjects). Statistical significance was tested with paired t-test (*p* = 0.18, 0.38, 0.03, 0.02 for gaps of 0.05, 0.1, 0.2, and 0.5 s, respectively, with Benjamini-Hochberg correction).

### Loudness and sound naturalness modulate facial responses

We then tested the sensitivity of ferret facial motion to natural sounds that carry different meanings or valence for the animals. We presented natural sound excerpts from five diverse categories (vocalizations from other species, natural environment, sounds from the animal facility, human speech/music, conspecific vocalizations) at three sound levels (65, 72, 80 dB). Time-averaged FME monotonically increased with sound level while sound category had no significant effect (Fig. 2A-C; significant effect of sound level: *p*_*loudness*_ < 10^*−*10^; non-significant effect of category: *p*_*category*_ = 0.62). This was confirmed by chance-level decoding of category identity from facial principal component statistics (SupFig. 3). To isolate sensitivity to higher-order natural structure (sound *naturalness* or behavioral meaning) while controlling for low-level acoustics, we generated synthetic versions of each natural excerpt matched in spectrotemporal statistics but lacking higher-order natural structure (Landemard et al., 2021). We predicted that natural sounds would elicit larger responses if recognized by the animals. We found no main effect of naturalness on FME, but we observed a significant interaction between naturalness and category (Fig. 2D, *p*_*naturalness*:*category*_ = 0.026). Paradoxically, for most categories FME was significantly larger for synthetic sounds, the opposite of what ecological salience would predict. This indicates that ferret facial responses distinguish natural from model-matched synthetic sounds in a category-dependent manner despite matched low-level acoustic structure, consistent with a heightened response to acoustically unfamiliar stimuli for specific sound textures.

**Figure 2.**
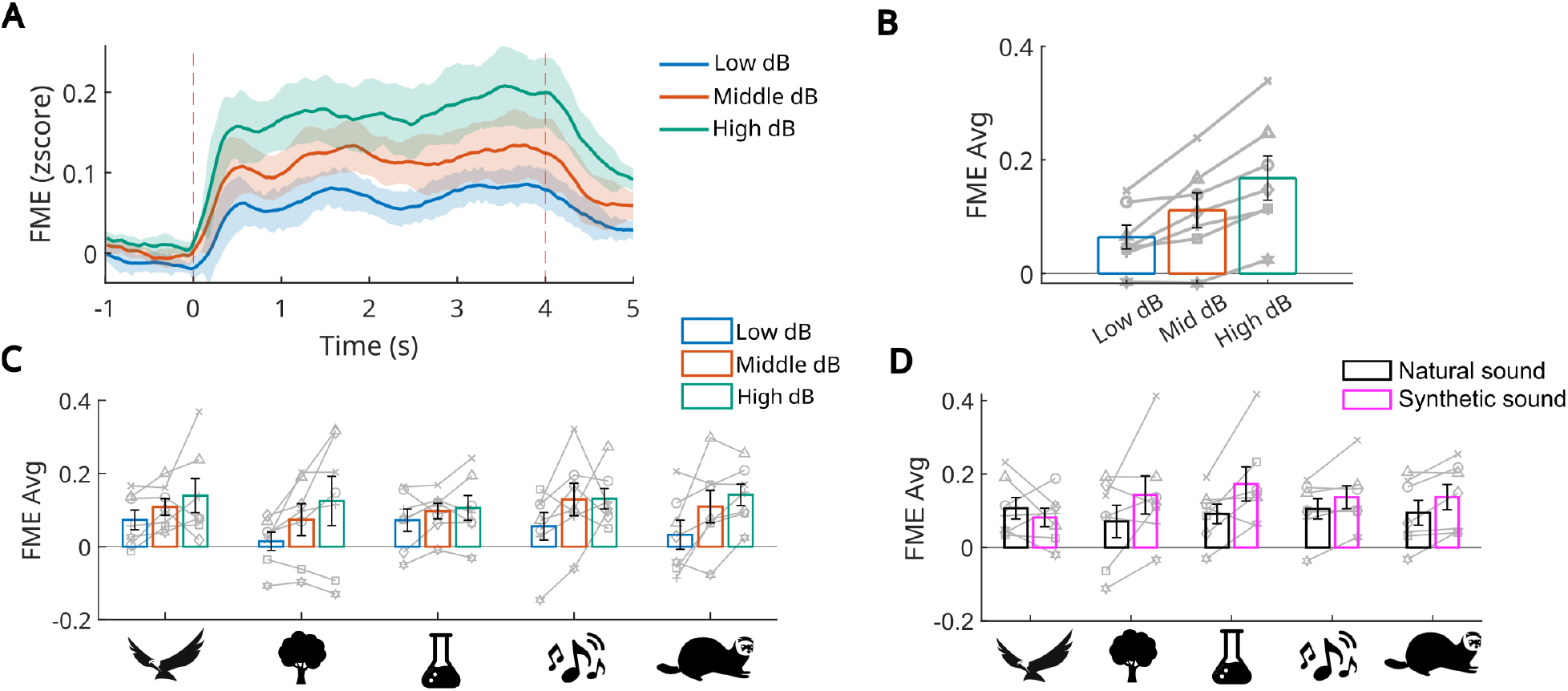
Loudness and sound naturalness modulate facial responses. **A.** Average FME responses across-subject and across sounds, for each sound level (low: 65 dB SPL, medium: 72 dB SPL, high: 80 dB SPL). **B**. Time-averaged FME as a function of sound level (gray lines: individual subjects). **C**. Time-averaged FME by sound level and sound category (gray lines: individual subjects). We observed a main effect of sound level but no effect of sound category (LME: sound-level effect: *F*_2,198_ = 28.78, *p*_*loudness*_ < 10^*−*10^; no category effect: *F*_4,198_ = 0.66, *p*_*category*_ = 0.62). **D**. Time-averaged FME comparing natural sounds with synthetic versions matched in spectrotemporal structure (gray lines: individual subjects). Naturalness showed no main effect (LME: *F*_1,198_ = 1.01, *p*_*naturalness*_ = 0.32) but significantly interacted with sound category (LME: *F*_4,198_ = 2.84, *p*_*naturalness*:*category*_ = 0.026).

### Facial motion responses reflect coarse but not fine temporal acoustic features

Although sound category was not represented in facial motion, it remained possible that FME tracked fine temporal features of individual sounds rather than a generic response to sound presence. To test this, we modeled FME as a linear function of four acoustic features using a temporal response function (TRF) encoding framework traditionally used in EEG (Crosse et al., 2016): a binary sound onset/offset regressor (called “On/Off” hereafter), a binarized envelope, a binarized envelope derivative, and the raw envelope. This enables the assessment of sensitivity to temporal events across different scales and dynamics, ranging from the sound presence to rapid acoustic transients. Models were trained using either single features, or combinations of multiple features (Fig. 3A), yielding both TRF kernels (Fig. 3B-D) and cross-validated FME predictions (Fig. 3E,F). Examples from individual subjects (Fig. 3B,C) illustrate inter-individual variability in TRF dynamics, with Ferret 3 showing a sharper response to sound onset, while Ferret 1 displayed a more sustained response profile. The On/Off regressor, which is in up state throughout sounds and then carries no stimulus-specific information, explained the largest fraction of FME variance in most animals (Fig. 3G,H). TRF weights of the On/Off regressor peaked at approximately 200 ms lag (Fig. 3B-D), consistent with the response latencies observed for broadband sounds (Fig. 1B-E). Additional acoustic features provided little improvement beyond the On/Off predictor alone (Fig. 3I), indicating that ferret facial motion reflects a generic response to sound presence rather than fine temporal sound structure. To ensure consistency, we validated that animals with larger facial movements exhibited higher prediction performance (SupFig. 4A) and that prediction accuracy for one sound feature correlated with that of another (SupFig. 4B). Together with the absence of category selectivity, these results suggest that sound-evoked facial motion in ferrets carries minimal stimulus-specific information, and therefore represents a behaviorally driven signal that can be largely characterized by a single onset-locked component.

**Figure 3.**
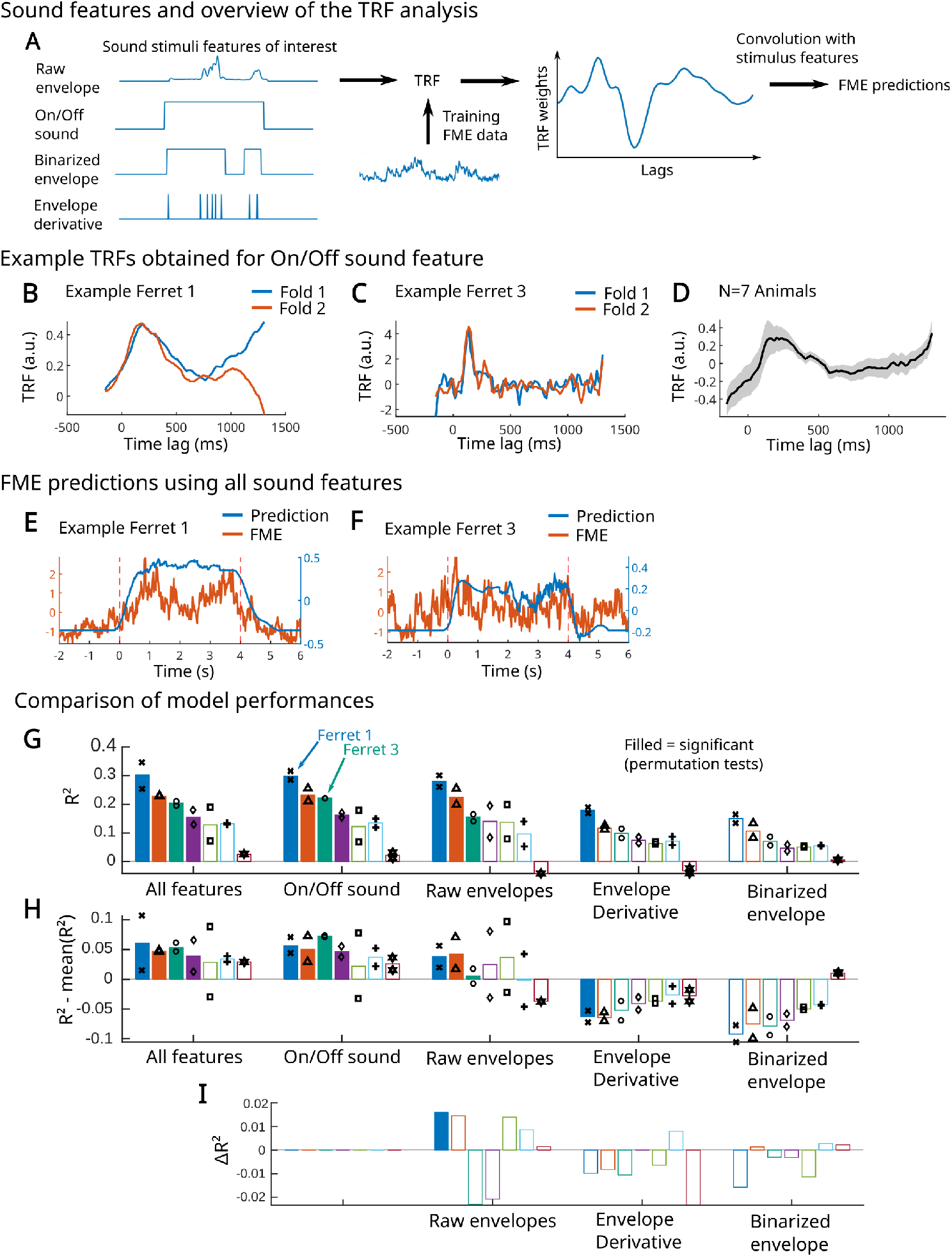
Facial responses reflects coarse sound temporal dynamics: temporal response function (TRF) analysis. **A.** Schematic of the TRF analysis (implemented using the mTRF toolbox (Crosse et al., 2016) and the sound features used as predictors: raw envelope, On/Off sound, binarized envelope, and envelope derivative. **B-C**. Example TRFs for single individuals when trained on the On/Off feature alone. **D**. Average TRF weights across animals when the model is trained on the On/Off feature alone. **E-F**. FME predictions for the same individual subjects when all features are combined in the model. **G**. Prediction R2 for each subject and each model trained on a single sound feature. Filled bars indicate statistically significant predictions (permutation test, *p* < 0.05, feature (Δ*R* = *R* Bonferroni corrected). **H**. Difference between each model’s R2 and the average R2 across models within the same animal. **I**. On/Off sound predominance (Δ*R*^2^ analysis): improvement in model performance when adding the x-axis sound feature to a model already including the On/Off sound feature 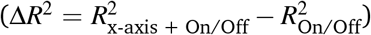. Filled bars indicate statistically significant improvement (permutation test, *p* < 0.05, Bonferroni corrected).

### Pupil dynamics mirror facial motion responses

We next asked whether pupil dynamics followed the same response pattern as facial motion, using Facemap-based pupil tracking (Syeda et al., 2024; Fig. 4A) as an independent measure of sound-evoked arousal changes. Overall, pupil responses were modulated by the same stimulus properties as facial motion. For natural sounds, pupil responses showed an initial transient constriction at sound onset, followed by pupil dilation (Fig. 4B), consistent with a sound-evoked change in arousal state. Pupil area increased with sound level and was larger for model-matched synthetic than natural sounds (Fig. 4B-C; significant effect of sound level: *p*_*loudness*_ = 0.013; significant effect of naturalness: *p*_*naturalness*_ < 10^*−*6^). Sound category showed no significant main effect (Fig. 4D; *p*_*category*_ = 0.68), but interacted significantly with naturalness (*p*_*naturalness*:*category*_ < 10^*−*2^), paralleling the FME results.

**Figure 4.**
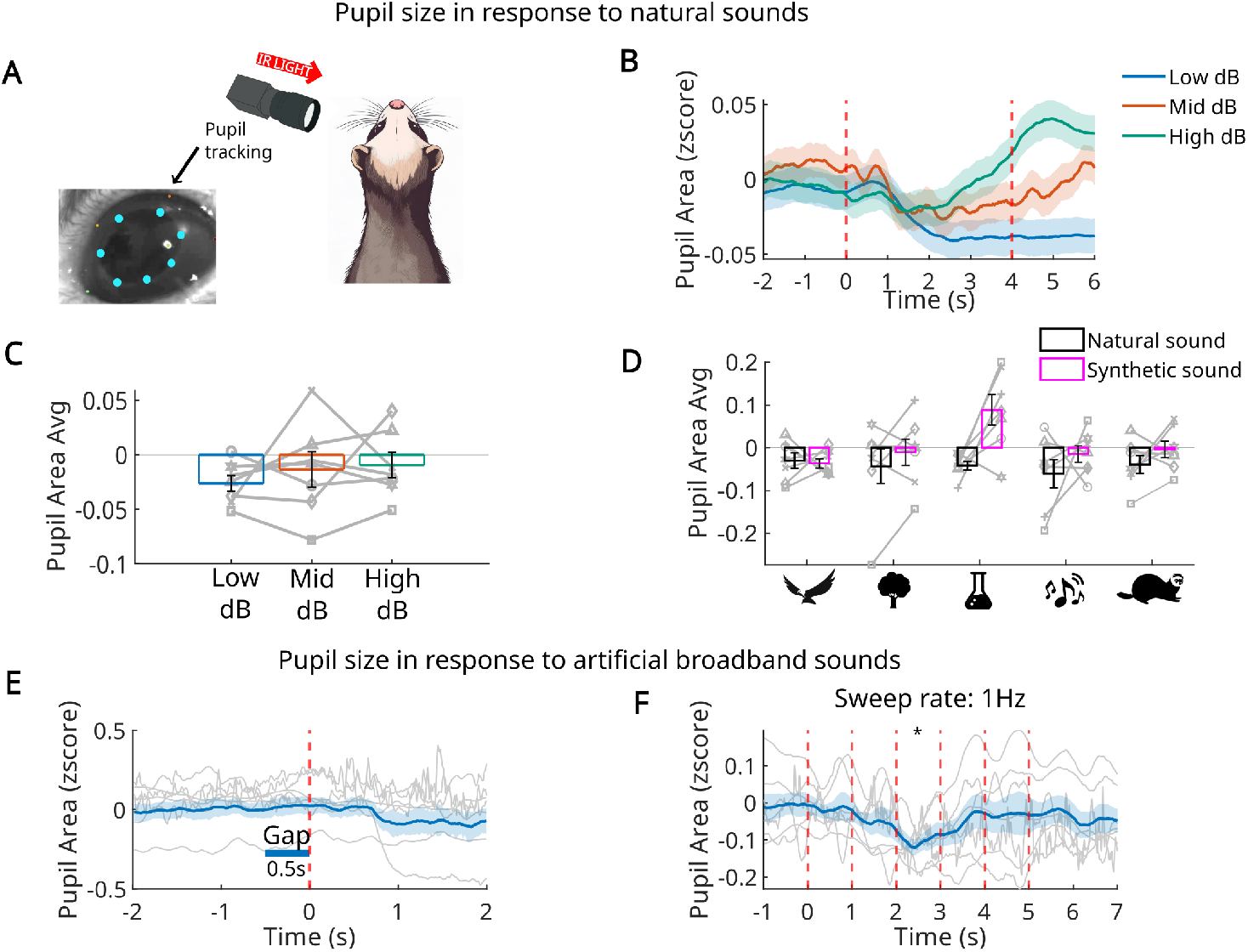
Pupil responses correlate with facial responses. **A.** Example of pupil tracking using Facemap (Syeda et al., 2024). **B**. Pupil area time courses averaged across all subjects and sounds, for each sound level. **C**. Time-averaged pupil area as a function of sound level (LME: significant effect of sound level: *F*_2,198_ = 4.40, *p*_*loudness*_ = 0.013, gray lines: individual subjects). **D**. Time-averaged pupil area as a function of sound naturalness (LME: significant effect of naturalness: *F*_1,198_ = 29.57, *p*_*naturalness*_ < 10^*−*6^; non-significant effect of category: *F*_4,198_ = 0.58, *p*_*category*_ = 0.68; significant interaction between naturalness and sound category: *F*_4,198_ = 4.12, *p*_*naturalness*:*category*_ < 10^*−*2^; gray lines: individual subjects). **E**. Pupil response to a 0.5-s silent gap (blue: across-subject mean; gray: individual subjects). Statistical significance was assessed using a paired t-test, the effect was not significant (*p* = 0.61) **F**. Pupil response to sweep trials at 1 Hz presentation rate (third sweep: *p* = 0.04, corrected paired t-test).

For broadband sounds, significant pupil constrictions were observed only for the slowest temporal structure, namely the slowest sweep (corrected paired t-test, Fig. 4F, p=0.04, SupFig. 5), while gap responses were not significant (Fig. 4E, SupFig. 5). This confirms that pupil dynamics, like facial motion, primarily reflect coarse rather than fine temporal sound structure.

## Discussion

Our results demonstrate that passive sounds reliably evoke uninstructed facial motion and pupil responses in head-fixed ferrets, establishing that sound-evoked behavioral responses are also present in carnivores. However, ferret responses differed from those described in mice in three key respects: they were slower, dominated by a generic onset-locked component rather than tracking fine temporal structure, and carried no information about sound identity or category. In mice, sound-evoked facial motion is robust, stereotyped, and sensitive to stimulus fine structure (Bimbard et al., 2023; Clayton et al., 2024; Musall et al., 2019). The present results suggest that ferrets occupy a distinct position along this spectrum, with sound-evoked movement present but less informative about stimulus content.

### Divergent sensorimotor coupling in carnivores and rodents

A central question motivating this study was whether ferrets, a carnivore species, fall along the continuum between mice, where stimuli evoke strong and rich movement that shape cortical responses, and primates, where spontaneous movements to stimuli are reduced, and contribute only weakly to neuronal activity in sensory cortices (Talluri et al., 2023). Our findings place ferrets closer to the primate end of this spectrum. Sound-evoked FME in ferrets was driven primarily by coarse acoustic features (sound presence and loudness) and did not track the rich temporal structure that characterizes facial responses in mice. In mice, facial motion synchronizes to sweep rates up to 3.5 Hz and responds reliably to gaps shorter than 0.1 s (Clayton et al., 2024); in ferrets, coherent synchronization was restricted to modulations below 2 Hz and only the longest gaps evoked significant responses. This extended temporal bandwidth in mice may underlie the greater stimulus specificity of their facial responses, which encode the fine temporal structure supporting sound discriminability lacking to ferret facial motion. Ferret responses were also less homogeneous across individuals than those reported in mice (Fig. 1, 3, 4; SupFig. 2, 5; (Bimbard et al., 2023; Clayton et al., 2024)), consistent with the view that tight coupling may reflect something specific to rodent sensorimotor organization rather than a general mammalian feature (Parker et al., 2020).

From a practical standpoint, this result is reassuring for ferret auditory neuroscience. Ferrets are an important non-rodent model in auditory research, valued for their large gyrified cortex, well-characterized auditory hierarchy, and processing properties routinely compared to those of humans (Landemard et al., 2021, 2025; Sabat et al., 2025). If sound-evoked facial motion in ferrets is dominated by a generic onset response that does not track stimulus-specific features, its contribution to cortical variability is likely more limited than in mice and more tractable to regress out. Nevertheless, the present data suggest that behavioral state and sound-onset responses are not fully negligible: monitoring facial motion and pupil dynamics alongside neural recordings remains advisable in ferret experiments, particularly when comparing conditions that differ in loudness or naturalness.

### Onset-driven behavioral response, arousal, and acoustic surprise

The 200–300 ms latency of the sound-evoked behavioral response is consistent with a motor response mediated by subcortical arousal circuits rather than cortical sensory encoding. The locus coeruleus–norepinephrine (LC-NE) system is a particularly strong candidate: LC neurons are phasically activated by unexpected or salient stimuli and discharge in tight temporal coupling with behavioral orienting responses, driving widespread network reorganization via noradrenaline release in forebrain targets (Sara & Bouret, 2012). Pupil dilation, widely used as a proxy for LC-NE activity, independently supports this interpretation: at high sound levels, the initial pupil constriction was followed by a sustained dilation phase consistent with a delayed noradrenergic arousal shift (Montes-Lourido et al., 2021; Zekveld et al., 2018). Like FME, pupil responses were restricted to the slowest temporal structures, further arguing for a common subcortical arousal mechanism.

This framework provides a natural account of the enhanced responses we observed to synthetic sounds. Natural sounds, by virtue of their higher-order statistical regularities, are more predictable given the animal’s prior auditory experience. Synthetic sounds, which preserve spectrotemporal statistics but destroy the higher-order structure of natural sound textures, may be perceived as acoustically anomalous, i.e., familiar in coarse spectral content but strange in their spectrotemporal organization. Within a predictive coding framework, such stimuli generate larger prediction errors, which would recruit the LC-NE system more strongly and produce the enhanced arousal and movement responses we observed. The fact that the synthetic-versus-natural effect was category-dependent is consistent with this view: only stimuli with sufficiently rich higher-order structure should generate detectable prediction errors at the level of facial motor output. An analogous phenomenon in human electrophysiology is the mismatch negativity (MMN), an automatic response to violations of auditory regularities that occurs even without directed attention (Näätänen et al., 2007). Whether ferrets exhibit an MMN-like response to higher-order statistical violations remains an interesting open question. This behavioral sensitivity also stands in interesting contrast to our own neuroimaging results in ferret auditory cortex: Landemard et al. (2021) showed that, unlike human auditory cortex, ferret auditory cortex does not strongly differentiate natural from spectrotemporally matched synthetic sounds at the mesoscopic level of neural population responses. One possibility is that this discrimination is encoded at a finer spatial or temporal scale than hemodynamic imaging can resolve, and that the behavioral response reflects a latent cortical sensitivity, motivating future electrophysiological studies of higher-order acoustic coding at the neuronal level.

### A somatotopic hypothesis for species differences in movement-neural coupling

The species differences documented here may reflect not merely quantitative differences in the degree of sensorimotor coupling, but a fundamental difference in which body part is most tightly coupled to sensory processing and behavioral state in each species. We propose that the body part occupying the largest cortical representation in a given species is also the one whose movements most strongly co-vary with arousal, and is therefore the most informative readout of sound-evoked behavioral changes. Support for this idea comes from within-species data: Mimica et al., 2023 found that movement encoding across rat neocortex is not uniform but body-part-specific, with different cortical areas preferentially representing different effectors, consistent with somatotopic organization extending well beyond classical sensorimotor regions. Extrapolating across species, it follows that the most informative movement signal for a given recording site would reflect the body part most heavily represented in the cortex, and that this relationship should vary predictably with the known somatotopic organization of each species.

In mice, the snout and whisker pad occupy a disproportionately large fraction of primary somatosensory cortex, organized into the barrel field in which each vibrissa maps onto a discrete cortical column (Petersen, 2007; Woolsey & Van der Loos, 1970). The face is the primary motor effector of sensory exploration in rodents, so it is unsurprising that facial motion is the most informative channel for predicting cortical state in that species. In primates, by contrast, hands and fingers dominate primary somatosensory and motor cortex alongside the face (Penfield & Rasmussen, 1950), reflecting the central role of dexterous manipulation in primate behavior. This hypothesis predicts that finger movements, not facial movements, should be the most informative uninstructed movement channel in primates. Consistent with this, Talluri et al. (2023) found that spontaneous movements contribute only weakly to primate visual cortex, but that study did not monitor fine finger movements. When hand movements were explicitly tracked, they emerged as among the most informative uninstructed variables accounting for neural variance in prefrontal cortex during task encoding (Tremblay et al., 2023), suggesting that the relevant movement channel had simply not been monitored in earlier work.

For ferrets, the forepaw has a substantial and well-organized representation in somatosensory cortex, occupying approximately the caudal half of the posterior sigmoid gyrus with multiple digit representations (McLaughlin et al., 1998). Under our experimental conditions, animals were head-fixed in a contention tube that substantially restricted forepaw movement, while only facial motion was monitored. If forepaw movements are the most informative channel in ferrets, this design would systematically underestimate movement-state coupling in this species. A direct test would involve simultaneously recording facial and forepaw motion alongside auditory cortical activity in freely moving ferrets, which would clarify whether the weak facial responses we observed reflect a genuine property of ferret sensorimotor organization or simply the choice of body part monitored. More broadly, a comparative cross-species study of movement-neural coupling across body parts would provide a powerful test of whether the mouse results reflect a general principle of sensorimotor integration or a rodent-specific specialization of the face as the primary interface between the animal and its sensory environment. Answering these questions has direct practical consequences for auditory neuroscience: understanding which movement channels contaminate cortical recordings in each species is a prerequisite for correctly interpreting what sound-evoked neural activity actually reflects.

## Acknowledgments

We thank Sophie Bagur for valuable discussion. This work was supported by ANR-17-EURE-0017 and ANR-10-IDEX-0001-02, PSL-NEURO, Fondation Pour l’Audition, Institut Universitaire de France, ANR-23-NEUC-0001 and ANR-CROCOS.

## Methods

### Experimental procedure

All experimental procedures were approved by the French Ministry of Agriculture (protocol authorization: 21022) and were conducted in full accordance with European legislation governing the protection of animals used for scientific purposes (2010/63/EU). Experiments were carried out on seven adult female ferrets. To ensure stability of the videos, animals were head-fixed while awake in a custom contention tube. A stainless-steel headpost was surgically implanted onto the skull under 1% isoflurane anesthesia (Chillale et al., 2023). More information about each animal’s history can be found in Supplementary Table 1. Sound stimuli were delivered passively in an open-field configuration inside a sound-isolated booth, with the animals positioned facing a speaker aligned to head height. Facial movements and pupil dynamics were simultaneously recorded. Facial video was acquired at 60 Hz with a monochromatic Basler camera equipped with a 25-mm focal length lens, while pupil diameter was monitored using a second monochromatic Basler camera at 80 Hz with an 8-mm focal length lens. The pupil was illuminated with an infrared LED (940 nm), supplemented by the booth’s inherent lighting.

### Stimuli

Broadband sounds stimuli and Natural sounds were presented in two separated experiments.

In the broadband sound experiments, two recording sessions (approximately 1 hour each) were collected for every animal. Each session contained 8 sequences presented in alternation: four sweep sequences and four gap sequences. Sequences were separated by a 30s pause. Sweeps and gaps trials were presented at 72dB SPL.

- **Sweep sequences:** Each sweep sequence consisted of 20 trials. Each trial contained six descending frequency sweeps (16 kHz → 500 Hz), each sweep lasting 250 ms. Sweeps were presented at one of four repetition rates (1, 2, 3, or 4 Hz). Within a sequence, each rate was used in five trials, in randomized order. The interval between trials was a uniformly randomized pause of 7-11 s.
- **Gap sequences:** Each gap sequence consisted of continuous broadband noise containing 20 silent gaps. The noise duration between gaps averaged 8 s and was randomized using a flat hazard function. Four gap durations were used (50, 100, 200, and 500 ms), each presented five times per sequence, in randomized order.

In the natural sound experiments, three recording sessions were conducted per ferret (each session consisting of two 1-hour recordings within a single day). Each session included four repetitions of every combination of sound identity and sound level, with the presentation order randomized. All sound excerpts lasted 4s and were followed by a randomly timed pause of 2 to 4 s. RMS-normalized sounds were calibrated to three sound levels: 65 dB SPL (low), 72 dB SPL (medium), and 80 dB SPL (high). 41 natural sounds were tested, selected from five categories of interest: other animals, natural environment, laboratory environment, human sounds, and ferret sounds (Table 1).

**Table 1.** Natural sound selected and their categories.

| Category | Other animals (predators and prey) | Natural environment | Laboratory environment | Human sounds (voices + music) | Ferret sounds |
| --- | --- | --- | --- | --- | --- |
| Source | Pro Sound Effects database | Pro Sound Effects database | Homemade recordings | Freely accessible diverse databases | Internal database |
|  | Owl 1 | Forest 1 | The laboratory animal caretaker's voice | French-speaking man | Fight call 1 |
|  | Owl 2 | Forest 2 | Feeding call 1 (kibbles) | French-speaking woman | Fight call 2 |
|  | Eagle 1 | River 1 | Feeding call 2 (feeder noise) | English-speaking man | Shock vocalization 1 |
|  | Eagle 2 | River 2 | Drinking call (water bottle) | English-speaking woman | Shock vocalization 2 |
|  | Mouse | Rain | Door opening | Salsa music | Pup call 1 |
|  | Frog 1 | Storm | Cage opening | Funk music | Pup call 2 |
|  | Frog 2 |  | Toy noise | Rock music | Pup call 3 |
|  | Blue tit 1 |  | Litter |  | Male vocalization 1 |
|  | Blue tit 2 |  | Tunnel |  | Male vocalization 2 |
|  |  |  | Background noise |  |  |

For each of these natural sounds, we synthesized a synthetic sound matching the cochlear statistics and spectrotemporal modulations. To do so, we used a custom algorithm described in Landemard et al., 2021; Norman-Haignere and McDermott, 2018. Briefly, cochleagrams were computed by filtering each sound waveform with a bank of pseudo-logarithmically spaced cochlear filters, followed by envelope compression to mimic cochlear amplification (McDermott & Simoncelli, 2011) . Spectral and temporal modulation statistics were extracted by further filtering cochleagrams using temporal ([1, 2, 8, 32] Hz) and spectral ([0.5, 1, 2, 4] cycles/octave) filters (Chi et al., 2005). Then spectrotemporal filters were generated as the outer product of all pairs of temporal and spectral filters in the 2D Fourier domain. Sounds were synthesized by iteratively modifying an initially unstructured noise stimulus to match the time-averaged statistics of filtered cochleagrams using a histogram-matching procedure.

Synthetic sounds were presented in the same manner as their corresponding natural sounds, in randomized order during the same sessions. This resulted in 82 distinctive sounds presented at three sound levels, with 12 repetitions of each unique combination per animal.

### Data Preprocessing and Analysis

Unless otherwise specified, all envelope traces shown in the figures represent the mean ± SEM.

### Facial motion energy (FME)

FME was computed as the frame-to-frame absolute difference in pixel intensity, averaged across all pixels in each frame. In addition, we extracted image-based components using Facemap (Syeda et al., 2024). For each animal, all video recordings within a given experiment were concatenated and decomposed jointly using singular value decomposition (SVD), yielding a single, experiment-specific SVD basis per animal (500 components in total, SupFig. 1-2).

### Pupil tracking

Pupil tracking was also performed using Facemap. For each subject, a set of key points outlining the pupil border was manually annotated and used to train a subject-specific tracking model. Pupil area was then computed from the tracked coordinates on a frame-by-frame basis.

All the following analyses were performed using Matlab2024b.

### Normalization

For the natural-sound recordings, FME, principal components and pupil time-series were z-scored using the mean and standard deviation calculated only during silent periods within each recording. For the broadband-sound recordings, z-scoring was performed using the mean and standard deviation computed over the entire recording.

### Facial synchronization

Facial synchronization (Fig. 1D) was quantified from facial motion energy (FME) signals in the frequency domain (Clayton et al., 2024). For each animal and sweep condition, FME responses were averaged across trials, and the fast Fourier transform was computed. Synchronization was defined as the power at the stimulus presentation frequency expressed in decibels relative to a high-frequency noise floor, estimated as the mean power across the highest frequency bins:

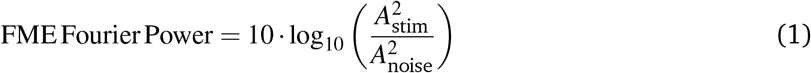

### Statistical tests

In Fig. 1, Fig. 4E-F, and SupFigs. 2 and 5, statistical significance of FME, principal components, and pupil-dilation responses to sweep trials and silent gaps was assessed using paired t-tests, comparing the response within a selected window to a 1-s baseline, averaged across all trials for each animal. For FME (Fig. 1, SupFig. 2), the response window was 0.35-0.50 s after noise onset for gap stimuli and 0.35-0.50 s after sweep onset for sweep stimuli, averaged across the first two sweeps to avoid adaptation in the following sweeps. For pupil area (Fig. 4, SupFig. 5), the response window was 1-2 s after noise onset for gap stimuli and 0.10-0.50 s after the third sweep onset for sweep stimuli, chosen to account for the pupil response lag. The Benjamini-Hochberg correction was applied to control for multiple comparisons across frequencies and gap durations.

In Fig. 1D, statistical significance was assessed using a repeated measures ANOVA performed on the average power measured at each sweep presentation rate for each animal, with presentation rate as the within-subject factor (model: ‘Rate1-Rate4 *∼* 1’).

In Fig. 2 and Fig. 4B-D, we assess the effects of stimuli properties on the time-averaged FME and pupil area using a linear mixed-effects model implemented in MATLAB (fitlme) with maximum likelihood (ML) estimation. The model included SoundLevel, Category and Naturalness (Natural vs Synthetic) as fixed effects, and a random intercept for Animal. After inspection of the data, we additionally included an interaction term between Naturalness and Category to test for category-dependent effects of naturalness: FMEAvg *∼* 1 +Naturalness *×* Category+SoundLevel+(1 | Animal). Fixed-effect significance was assessed using t-tests.

### Temporal Response Function analysis

To assess how facial motion energy (FME) evolves in response to natural sounds, and specifically to determine which sound features and their temporal dynamics are most strongly encoded in FME, we adapted a modelling framework commonly used in EEG data analysis: the temporal response function (TRF) approach, implemented using the mTRF-Toolbox (Crosse et al., 2016).

In our implementation we used a forward (encoding) model: stimulus features (here, sound-specific feature time-courses) were convolved with a linear filter (the TRF) to predict the target outcome (FME here, rather than neural activity, as in the original applications of the toolbox). The toolbox estimates the filter coefficients via regularised ridge regression, across a specified range of time-lags between stimulus and outcome. This procedure captures how changes in the stimulus features map over time into the predicted variable. Once the optimal regularisation parameter is selected using cross-validation, the resulting TRF coefficients can be interpreted as the temporal weighting of each feature at each lag (i.e., how strongly and at what latency a unit change in the stimulus feature contributes to predicted FME).

We applied the model to four distinct sound features, chosen to represent different temporal dynamics:

- **Raw envelope:** absolute value of the Hilbert transform of the sound waveform, smoothed.
- **Binary onset time-course:** value = 1 throughout sound presentation, 0 during silence.
- **Binarised envelope:** applying a fixed threshold to the raw envelope, values below threshold = 0, above = 1.
- **Impulsed derivative of the envelope:** applying a fixed threshold to the derivative of the envelope, values below threshold = 0, above = 1.

Models were trained either on each isolated feature or on combinations of features. In the manuscript we report primarily the isolated-feature results (except for the combined-feature prediction shown in Fig. 3E-F and Δ*R*^2^ analysis Fig. 3I), and we trained models for each animal individually.

Prior to model fitting, FME signals were averaged across identical trials (same sound identity and sound level), excluding low sound level conditions (65 dB), in order to improve signal-to-noise ratio and isolate stimulus-driven responses.

For each model, we first conducted a two-fold cross-validation to select the optimal ridge regularization parameter. Then, with that parameter fixed, we conducted a second two-fold cross-validation to estimate the TRFs and generate predicted FME time-courses. We used a time-lag window from t_*min*_ = -150 ms to t_*max*_= +1300 ms.

Statistical significance of the prediction performance was assessed via a permutation test (n = 1000 permutations). For each permutation the correspondence between stimulus feature and FME was randomly reassigned (while keeping the same ridge parameter as before) and we then estimated the TRFs and the predictions (cross-validated). A significance threshold of *α* = 0.05 was applied, with correction for multiple comparisons across subjects and sound features using a Bonferroni correction.

For the Δ*R*^2^ analysis, we selected the feature identified as most informative in Fig. 3G and 3H, namely the On/Off sound feature. We first computed a baseline model including only the On/Off predictor and obtained its corresponding *R*^2^. We then added each additional feature separately to this baseline model and recomputed *R*^2^. The incremental contribution of each feature was quantified as Δ*R*^2^, defined as the difference in explained variance between the extended and baseline models 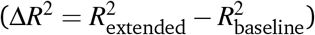. Statistical significance of the observed Δ*R*^2^ values was assessed using a permutation test (n=200 permutations). To account for multiple comparisons across features, a Bonferroni correction was applied.

### Classifier and t-SNE

Facial motion features were extracted from the first 50 singular value decomposition (SVD) components (SupFig. 3). For each sound and sound level, SVD features (means and variances) were averaged across identical trials, excluding low sound level conditions (65 dB). Classification was performed separately for each individual using a linear discriminant analysis (LDA) classifier with leave-one-out cross-validation. Performance was quantified using overall accuracy and confusion matrices. Statistical significance was assessed using permutation tests (n = 1000 permutations) and corrected for multiple comparisons across individuals using the Benjamini-Hochberg procedure. To visualize the structure of the feature space, t-distributed stochastic neighbor embedding (t-SNE) was applied to the feature vectors (2 dimensions, perplexity = 30), color-coded by sound category.

## Supplementary Figures

**Supplementary Figure 1:**
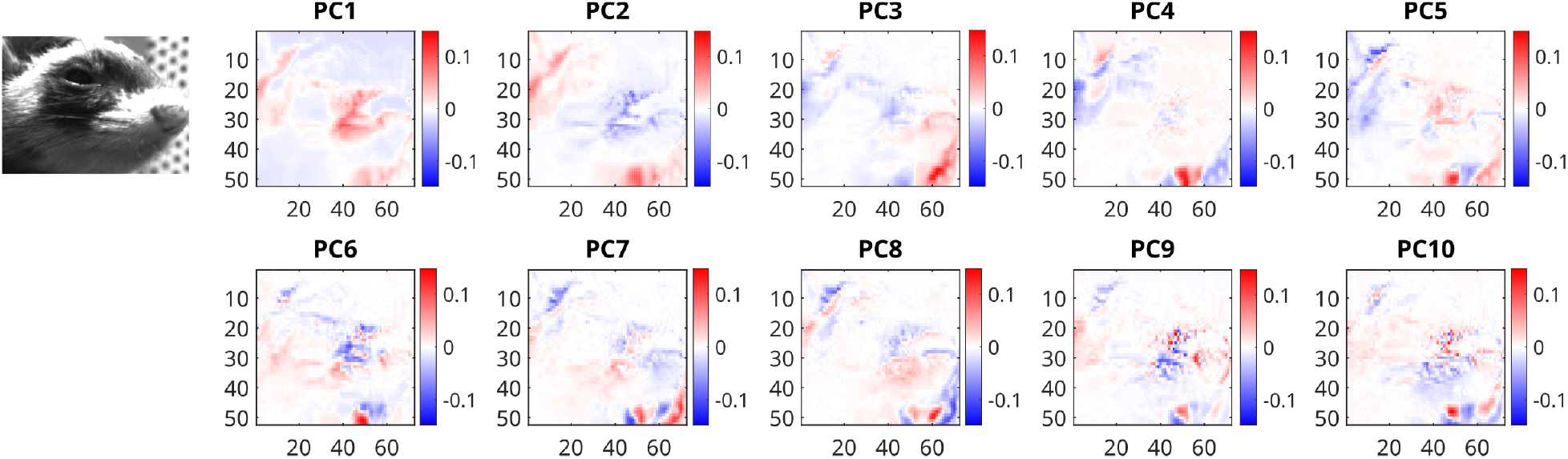
Single-individual example of the first ten motion components (masks) extracted via FaceMap (Syeda et al., 2024).

**Supplementary Figure 2:**
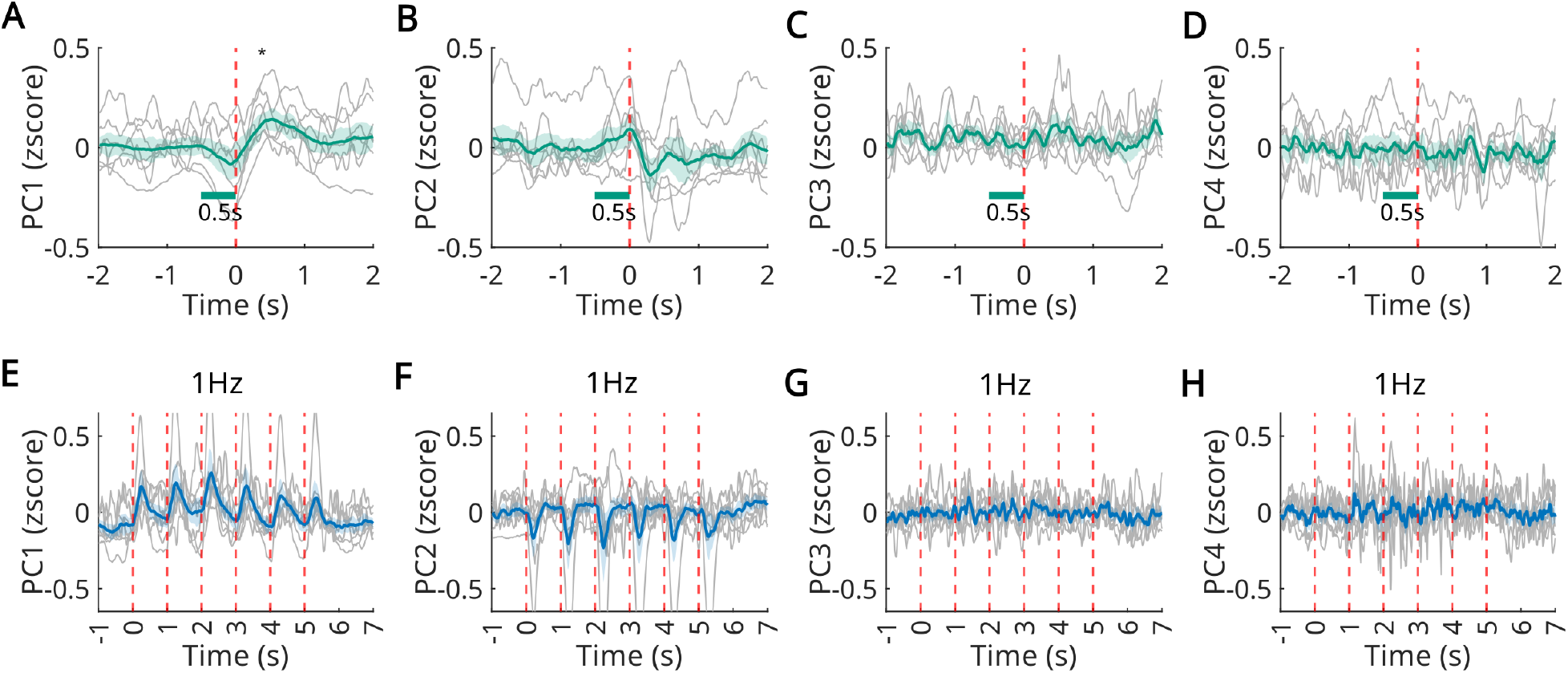
Principal component responses to broadband sounds. **A-D**. Responses of principal components 1-4 to a 0.5 s silent gap embedded in broadband noise. Statistical significance was assessed with paired t-tests (for the 0.5 s gap: *p* = 0.01, 0.27, 0.58, and 0.73 for PC1-4, respectively), with Benjamini-Hochberg correction for multiple comparisons across the four gap durations for each PC. **E-H**. Responses to descending frequency sweeps presented at a rate of 1 Hz. The sweep-evoked response was tested with a paired t-test on the mean response to the first two sweeps (at 1Hz: *p* = 0.03, 0.93, 0.07, and 0.94 for PC1-4, respectively), with Benjamini-Hochberg correction for multiple comparisons across the four sweep rates for each PC.

**Supplementary Figure 3:**
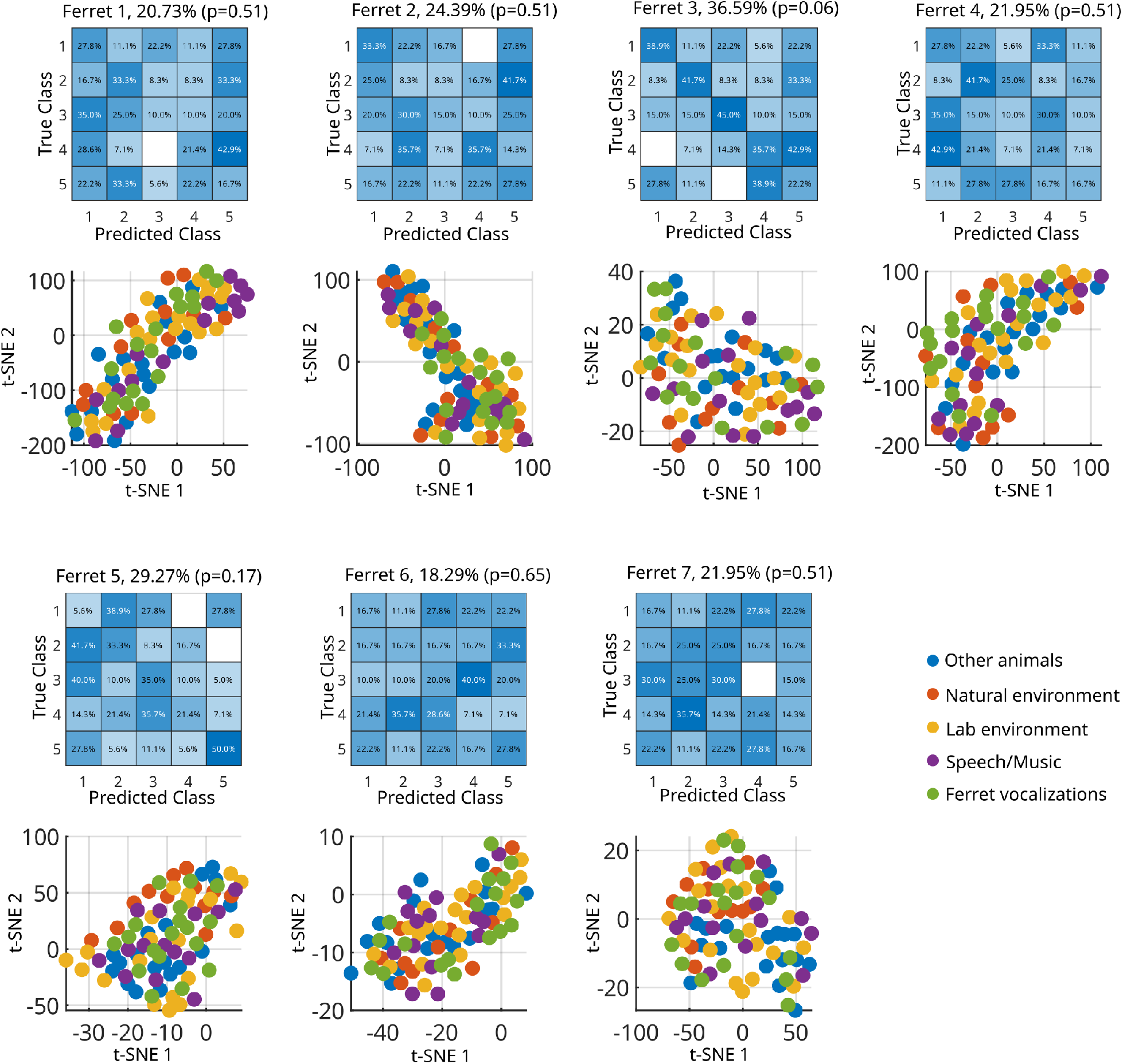
Decoding sound categories using statistics of the first 50 SVD components. For each ferret, the top panels show confusion matrices of a linear discriminant analysis (LDA) classifier, accuracy in percentages and statistical significance assessed via permutation tests and Benjamini-Hochberg correction. The bottom panels display t-SNE embeddings of the SVD statistics used as inputs to each classifier. Each dot represents data corresponding to a single sound at a given sound level, averaged across all identical trials.

**Supplementary Figure 4:**
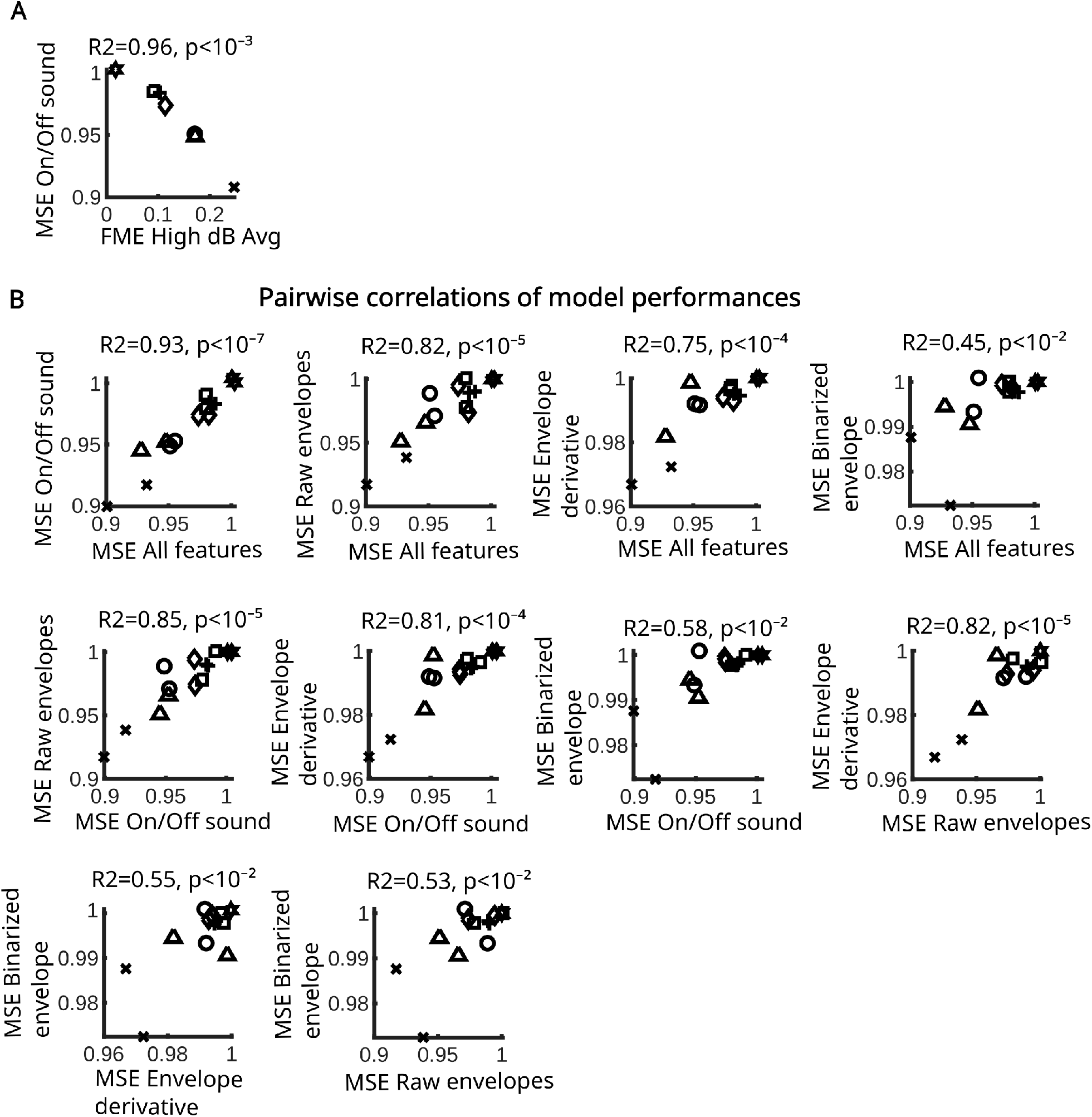
Additional analyses of TRF models performances A. MSE of the On/Off sound model versus the average FME amplitude at high sound level. **B**. Pairwise correlations of model performances.

**Supplementary Figure 5:**
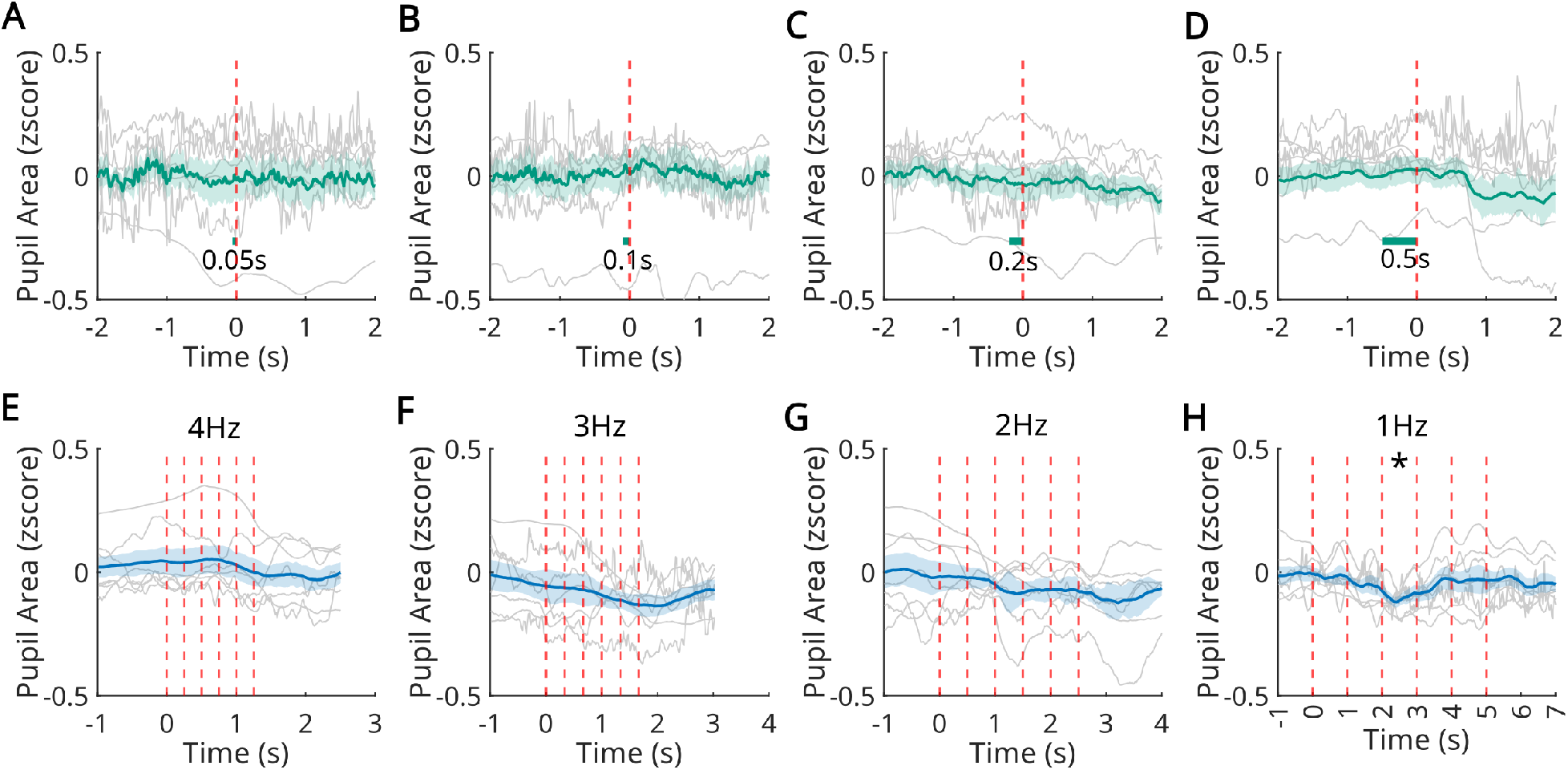
Pupil size responses to broadband sounds. **A-D**. Responses of pupil size to silent gaps embedded in broadband noise with durations of 0.05 s, 0.1 s, 0.2 s, and 0.5 s, respectively. Statistical significance was tested with paired t-test (*p* = 0.61, 0.76, 0.22, 0.61 for gaps of 0.05, 0.1, 0.2, and 0.5 s, respectively, with Benjamini-Hochberg correction for multiple comparisons). **E-H**. Responses to descending frequency sweeps presented at repetition rates of 4 Hz, 3 Hz, 2 Hz, and 1 Hz, respectively. Statistical significance was tested with paired t-tests on the response to the third sweep (*p* = 0.04, 0.49, 0.14, 0.44 for sweep rates 1-4 Hz), Benjamini-Hochberg corrected for multiple comparisons.

**Supplementary Table 1.**
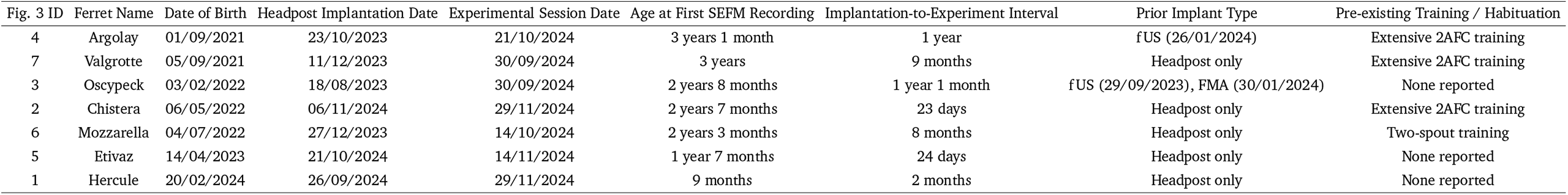
Summary of subject histories.

## References

Bimbard, C., Sit, T. P. H., Lebedeva, A., Reddy, C. B., Harris, K. D., & Carandini, M. (2023). Behavioral origin of sound-evoked activity in mouse visual cortex [Publisher: Nature Publishing Group]. Nature Neuroscience, 26(2), 251–258. 10.1038/s41593-022-01227-x

Chi, T., Ru, P., & Shamma, S. A. (2005). Multiresolution spectrotemporal analysis of complex sounds. The Journal of the Acoustical Society of America, 118(2), 887–906. 10.1121/1.1945807

Chillale, R. K., Shamma, S., Ostojic, S., & Boubenec, Y. (2023). Dynamics and maintenance of categorical responses in primary auditory cortex during task engagement (A. J. King, Ed.). eLife, 12, e85706. 10.7554/eLife.85706

Clayton, K. K., Stecyk, K. S., Guo, A. A., Chambers, A. R., Chen, K., Hancock, K. E., & Polley, D. B. (2024). Sound elicits stereotyped facial movements that provide a sensitive index of hearing abilities in mice [Publisher: Elsevier]. Current Biology, 34(8), 1605–1620.e5. 10.1016/j.cub.2024.02.057

Crosse, M. J., Di Liberto, G. M., Bednar, A., & Lalor, E. C. (2016). The multivariate temporal response function (mTRF) toolbox: A MATLAB toolbox for relating neural signals to continuous stimuli [Publisher: Frontiers]. Frontiers in Human Neuroscience, 10. 10.3389/fnhum.2016.00604

Landemard, A., Bimbard, C., & Boubenec, Y. (2025). Hierarchical encoding of natural sound mixtures in ferret auditory cortex (A. J. King, Ed.). eLife, 14, RP106628. 10.7554/eLife.106628

Landemard, A., Bimbard, C., Demené, C., Shamma, S., Norman-Haignere, S., & Boubenec, Y. (2021). Distinct higher-order representations of natural sounds in human and ferret auditory cortex (J. M. Groh, A. J. King, G. Cogan, & T. Overath, Eds.) [Publisher: eLife Sciences Publications, Ltd]. eLife, 10, e65566. 10.7554/eLife.65566

McDermott, J. H., & Simoncelli, E. P. (2011). Sound Texture Perception via Statistics of the Auditory Periphery: Evidence from Sound Synthesis. Neuron, 71(5), 926–940. 10.1016/j.neuron.2011.06.032

McLaughlin, D. F., Sonty, R. V., & Juliano, S. L. (1998). Organization of the forepaw representation in ferret somatosensory cortex. Somatosensory & Motor Research, 15(4), 253–268. 10.1080/08990229870673

Meyer, A. F., Poort, J., O’Keefe, J., Sahani, M., & Linden, J. F. (2018). A head-mounted camera system integrates detailed behavioral monitoring with multichannel electrophysiology in freely moving mice. Neuron, 100(1), 46–60.e7. 10.1016/j.neuron.2018.09.020

Mimica, B., Tombaz, T., Battistin, C., Fuglstad, J. G., Dunn, B. A., & Whitlock, J. R. (2023). Behavioral decomposition reveals rich encoding structure employed across neocortex in rats. Nature Communications, 14(1), 3947. 10.1038/s41467-023-39520-3

Montes-Lourido, P., Kar, M., Kumbam, I., & Sadagopan, S. (2021). Pupillometry as a reliable metric of auditory detection and discrimination across diverse stimulus paradigms in animal models [Publisher: Nature Publishing Group]. Scientific Reports, 11(1), 3108. 10.1038/s41598-021-82340-y

Musall, S., Kaufman, M. T., Juavinett, A. L., Gluf, S., & Churchland, A. K. (2019). Single-trial neural dynamics are dominated by richly varied movements. Nature neuroscience, 22(10), 1677–1686. 10.1038/s41593-019-0502-4

Näätänen, R., Paavilainen, P., Rinne, T., & Alho, K. (2007). The mismatch negativity (MMN) in basic research of central auditory processing: A review. Clinical Neurophysiology, 118(12), 2544–2590. 10.1016/j.clinph.2007.04.026

Niell, C. M., & Stryker, M. P. (2010). Modulation of visual responses by behavioral state in mouse visual cortex [Publisher: Elsevier]. Neuron, 65(4), 472–479. 10.1016/j.neuron.2010.01.033

Norman-Haignere, S. V., & McDermott, J. H. (2018). Neural responses to natural and model-matched stimuli reveal distinct computations in primary and nonprimary auditory cortex [Publisher: Public Library of Science]. PLOS Biology, 16(12), e2005127. 10.1371/journal.pbio.2005127

Olsen, T., & Hasenstaub, A. (2025). Sensory origin of visually evoked activity in auditory cortex. Cell Reports, 44(10), 116364. 10.1016/j.celrep.2025.116364

Parker, P. R. L., Brown, M. A., Smear, M. C., & Niell, C. M. (2020). Movement-related signals in sensory areas: Roles in natural behavior. Trends in Neurosciences, 43(8), 581–595. 10.1016/j.tins.2020.05.005

Penfield, W., & Rasmussen, T. (1950). The cerebral cortex of man; a clinical study of localization of function [Pages: xv, 248]. Macmillan.

Petersen, C. C. H. (2007). The functional organization of the barrel cortex. Neuron, 56(2), 339–355. 10.1016/j.neuron.2007.09.017

Sabat, M., Gouyette, H., Gaucher, Q., Espejo, M. L., David, S. V., Norman-Haignere, S., & Boubenec, Y. (2025). Neurons in auditory cortex integrate information within a constrained and context-invariant temporal window. Current Biology, 35(24), 6114–6125.e7. 10.1016/j.cub.2025.11.011

Sara, S. J., & Bouret, S. (2012). Orienting and reorienting: The locus coeruleus mediates cognition through arousal. Neuron, 76(1), 130–141. 10.1016/j.neuron.2012.09.011

Stringer, C., Pachitariu, M., Steinmetz, N., Reddy, C. B., Carandini, M., & Harris, K. D. (2019). Spontaneous behaviors drive multidimensional, brainwide activity [Publisher: American Association for the Advancement of Science]. Science, 364(6437), eaav7893. 10.1126/science.aav7893

Syeda, A., Zhong, L., Tung, R., Long, W., Pachitariu, M., & Stringer, C. (2024). Facemap: A framework for modeling neural activity based on orofacial tracking [Publisher: Nature Publishing Group]. Nature Neuroscience, 27(1), 187–195. 10.1038/s41593-023-01490-6

Talluri, B. C., Kang, I., Lazere, A., Quinn, K. R., Kaliss, N., Yates, J. L., Butts, D. A., & Nienborg, H. (2023). Activity in primate visual cortex is minimally driven by spontaneous movements [Publisher: Nature Publishing Group]. Nature Neuroscience, 26(11), 1953–1959. 10.1038/s41593-023-01459-5

Tremblay, S., Testard, C., DiTullio, R. W., Inchauspé, J., & Petrides, M. (2023). Neural cognitive signals during spontaneous movements in the macaque. Nature Neuroscience, 26(2), 295–305. 10.1038/s41593-022-01220-4

Woolsey, T. A., & Van der Loos, H. (1970). The structural organization of layer IV in the somatosensory region (S I) of mouse cerebral cortex: The description of a cortical field composed of discrete cytoarchitectonic units. Brain Research, 17(2), 205–242. 10.1016/0006-8993(70)90079-X

Yeomans, J. S., Li, L., Scott, B. W., & Frankland, P. W. (2002). Tactile, acoustic and vestibular systems sum to elicit the startle reflex. Neuroscience & Biobehavioral Reviews, 26(1), 1–11. 10.1016/S0149-7634(01)00057-4

Yin, C., Melin, M. D., Rojas-Bowe, G., Sun, X. R., Couto, J., Gluf, S., Kostiuk, A., Musall, S., & Churchland, A. K. (2025). Spontaneous movements and their relationship to neural activity fluctuate with latent engagement states. Neuron, 113(18), 3048–3063.e5. 10.1016/j.neuron.2025.06.001

Zekveld, A. A., Koelewijn, T., & Kramer, S. E. (2018). The pupil dilation response to auditory stimuli: Current state of knowledge. Trends in Hearing, 22, 2331216518777174. 10.1177/2331216518777174

